# Evolutionary rescue is determined by differential selection on demographic rates and density dependence

**DOI:** 10.1101/740365

**Authors:** Anna C Vinton, David A Vasseur

## Abstract

Accelerated rates of climate change are expected to either lead to populations adapting and persisting, or suffering extinction. Traditionally ecological models make extinction predictions based on how environmental change alters the intrinsic growth rate (*r*). However, these often ignore potential for evolutionary rescue, or to avoid extinction via adaptive evolution. Moreover, the environment may impose selective pressure on specific demographic rates (birth and death) rather than directly on r (the difference between the birth and death rates). Therefore, when we consider the potential for evolutionary rescue, populations with the same r can have different abilities to persist amidst environmental change. We can’t adequately understand evolutionary rescue without accounting for demography, and interactions between density dependence and environmental change. Using stochastic birth-death population models, we found evolutionary rescue more likely when environmental change alters birth rather than the death rate. Furthermore, species that evolve via density dependent selection are less vulnerable to extinction than species that undergo selection independent of population density. Resolving the key demographic factors affected by environmental change can lead to an understanding of how populations evolve to avoid extinction. By incorporating these considerations into our models we can better predict how species will respond to climate change.

## Introduction

Environmental change can lead to a decrease in population growth rate, resulting in extinction in some cases, and persistence in others. The ability of a population to rebound by attaining a positive growth rate following environmental change is ultimately what allows it to avoid extinction. The need to understand the mechanisms underlying population rebound has spurred studies about demographic rescue (via immigration) and genetic rescue (via an increase in genetic variation) (Brown and Kodric-Brown 1977; Hufbauer et al. 2015; Whiteley et al. 2015). More recently evolutionary rescue, or population rebound due to an increase in density of an adaptive genotype, is a potential mechanism, operating in organisms ranging from microbes (Bell and Gonzalez 2009; Zhang and Buckling 2011) to insects (Agashe et al. 2011) and mammals (Mills et al. 2018).

The search for what makes evolutionary rescue probable has led to an increasing effort to find experimental, empirical and theoretical evidence (Gomulkiewicz and Holt 1995; Orr and Unckless 2008; Bell and Gonzalez 2009, 2011; Johannesson et al. 2011; Gonzalez et al. 2013; Lindsey et al. 2013; Martin et al. 2013; Ramsayer et al. 2013; Mills et al. 2018). Four primary factors affect the propensity for evolutionary rescue (Bell and Gonzalez 2009): as initial population size (Ramsayer et al. 2013), genetic variability due to standing genetic variation and mutations (Orr and Unckless 2008), genetic variability due to dispersal (Mills et al. 2018), and the extent and severity of environmental change (Lindsey et al. 2013). Although these results have advanced our understanding of how population growth rate can increase following decline due to environmental change, we still lack a clear understanding of the role of the underlying demographic rates. This is in part because there is wide variation in how environmental change alters population demographic rates (birth and death rates) that is not always explicitly represented in our model frameworks.

The environment can reduce population growth rate by decreasing the birth rate, increasing the death rate, or some combination of the two (Dempster 1983; Mccredie et al. 1983; Aanes et al. 2000; Sibly et al. 2000, 2005; Clutton-Brock and Coulson 2002; Crump et al. 2004; Brewer and Peltzer 2009). To generalize across taxa, previous studies investigating evolutionary rescue commonly model demographic rates using deterministic models that do not differentiate how the environment acts on the birth and death rates, but rather use a fixed parameter, the intrinsic rate of population increase, *r* (the difference between the birth rate and the death rate). Consequently, information about changes in a particular demographic rate can be lost if *r* is the focus of a study.

Populations with the same *r*, but different underlying demographic rates, may respond quite differently to environmental selection, affecting how quickly and effectively they adapt (Holt 1990). Take the case of two populations, where one has a high birth and death rate, while another has a low birth and death rate. If the difference between the two rates is equal, both populations will have the same *r*. But, all else held equal, the population with the higher birth and death rate will have a faster rate of population turnover, and will evolve in response to selection more quickly than the population with the low birth and death rate. A logistic or exponential growth model that depends on a single *r* value, doesn’t allow exploration of how selection and environment affect birth and death rates, and ultimately population extinction or persistence. Therefore, treating birth and death rates explicitly can give insight into which natural populations are more likely to persist via evolutionary rescue in the face of environmental change. So, the potential for successful evolutionary rescue of small populations depends explicitly on birth and death rates, not *r*, which abstracts away from these rates and obscures the actual speed of adaptation by ignoring the rate of population turnover.

Density dependence can also affect birth and death rates, and has been shown to influence the dynamics of many species (Sibly et al. 2000; Coulson et al. 2001; Reed and Slade 2008; Ouyang et al. 2014). Environmental change may or may not alter the strength of density dependence; this varies across taxa and type of change (Owen-smith 1990; Sibly et al. 2000; Coulson et al. 2001). For example, environmental change leading to a drought may decrease the availability or accessibility of resources (Owen-smith 1990), intensifying density dependence as individuals compete for water-limited resources. Theoretical studies predict that compensatory density dependence, or decrease in growth rate at high densities and increase at low densities, would allow for a larger population size following environmental change (Holt 1990; Ferguson and Ponciano 2015), further facilitating adaptation to new environments. Therefore, establishing the interaction between density dependence and environmental change in different demographic rates is of the utmost importance as the population size following an environmental perturbation determines the probability of extinction.

In the past, determining how the environment alters demographic rates in a way that is the most mathematically simple has been sufficient, but, we argue is no longer sufficient when our interest turns to persistence via evolutionary rescue. Therefore, initial studies of evolutionary rescue focusing on *r* for simplicity, need to be expanded because: (i) they underlying demographic parameters, and (ii) the interaction between environmental change and density dependence may strongly affect evolution. We investigate how the evolutionary rescue depends in detailed ways on how environmental change affects population demographic rates. Here we incorporate environmental conditions and their effects on density dependence into per-capita rates of birth and death, to elucidate their effect on population dynamics and persistence in a stochastic model. We find that populations where the environment affects their death rate as opposed to their birth rate are the most vulnerable to extinction. Furthermore, when environmental change intensifies density dependence, populations are better able to rebound from small population sizes and undergo evolutionary rescue.

## Methods

### Model Formulation

We construct a continuous-time individual-based logistic growth model, then consider four ways that environmental change might alter population demographic rates. In all cases, as these are logistic growth models, either the birth or death rate is density dependent. In Cases 1a and 1b, the environment alters the birth rate in a density independent and density dependent manner respectively. In Cases 2a and 2b, the environment alters the death rate similarly, in a density independent, and density dependent way. We pay particular attention to ensuring that the four cases converge on the same outcome when the environment is static, to best isolate the effects of life-history and selection on evolutionary rescue.

### Logistic growth

All of our model cases are rooted in the logistic growth equation where *g*(*N*) is a function describing the density dependence of the per-capita growth rate. Note that *g*(*N*) = *r* in the case of exponential growth, as there is no density dependence. We begin with a general equation for the rate of population growth,

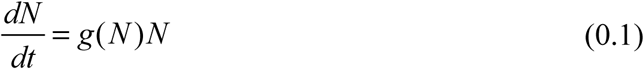

Since *g*(*N*) is equal to the difference between the per capita birth rate *b* and the per capita death rate *d*, we can rewrite equation (0.1) as the difference between birth and death rate,

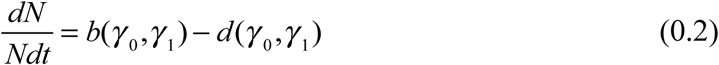

given that both birth and death are governed by density-independent *γ*_0_, and density-dependent *γ*_1_ contributions. Consistent with most derivations of logistic growth (Nåsell 1996, 2001), we assume that density-dependent factors tend to reduce birth rates and increase death rates, leading to the following general definition for birth and death rate functions:

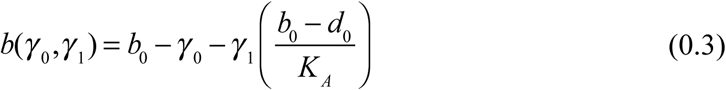

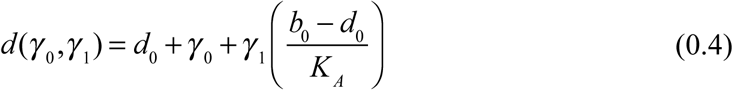

Where *b*_0_, *d*_0_ represents a modification of the background density-independent birth and death rates and *K*_*A*_ is the maximum carrying capacity.

### Environmental effect

We allow different demographic rates to depend on the environment and traits of individuals. For simplicity and tractability we model the environment *μ*_0_ as a simple sinusoidal function of time (see discussion for our reasons for this choice)

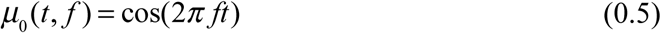

where *f* is frequency. We allow individuals to exhibit varied responses to the environment depending on their trait value *μ*. The effect of the environment, modulated by the trait, is given by

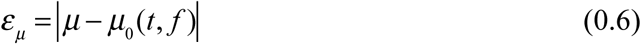

where a large *ε*_*μ*_ represents a maladapted individual, and a small *ε*_*μ*_ represents a well-adapted individual. We systematically incorporate the environmental effect *ε*_*μ*_ into the density independent *γ*_0_ and density dependent *γ*_1_ components of the birth and death rates. However, to facilitate comparison among the model cases, we scale our equations so that for any value of *ε*_*μ*_, the equilibrium population size (assuming no temporal environmental change) is the same across all of the model cases. This allows us to make an exact comparison of the impact of temporal environmental change on population dynamics, mediated by ecology and evolution. We do this by assuming that K_A_ represents the carrying capacity when an entire population is perfectly-adapted to their environment (*ε*_*μ*_ = 0) and we introduce a second carrying capacity, K_B_ for a population that is maladapted to their environment (*ε*_*μ*_ = 2). We then independently solve the parameters *γ* _0_ and *γ* _1_ given the conditions for carrying capacity. When the environment enters via a density-independent route (Cases 1a, 2a), we find:

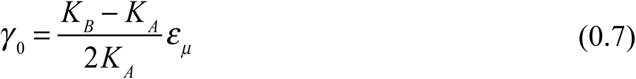

and *γ*_1_ for the models where the density dependence is altered by environmental change (Cases 1b, 2b)

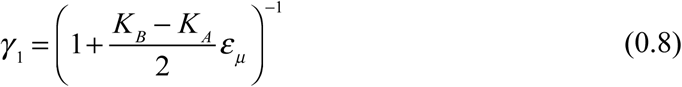

### Cases 1a-1b, dynamic birth models

We begin with Case 1a, where the environment alters *b*(*γ*_0_, *γ*_1_) in a density independent way as shown in Figure 1a. For case 1b, the environment again alters *b*(*γ*_0_, *γ*_1_) but in this case, it alters population response to density as shown in Figure 1b. For both dynamic birth models we hold *d*(*γ*_0_, *γ*_1_) constant and equal to *d*_0_.

**Figures 1 a-d.**
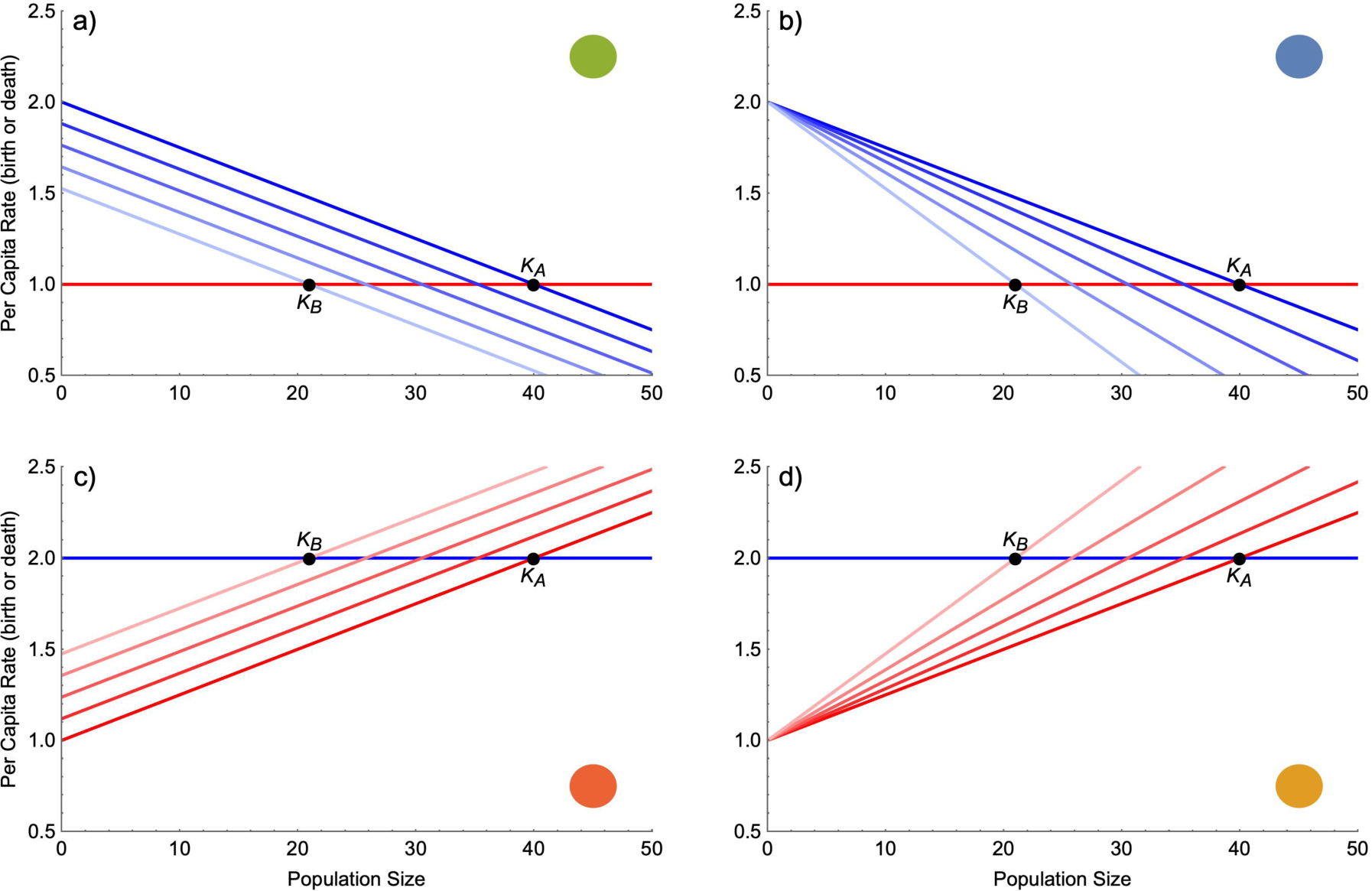
All four cases yield logistic population growth, but depend on different relationships between per-capita demographic rates of birth (blue) and death (red), (see equations 1.3 and 1.4, and table 1). In the upper panels (cases 1a, 1b) death rate is constant, birth rate is density dependent, and the environment either directly increases or decreases birth rate (1a) or changes the strength of the relationship between density and birth rate (i.e., density dependence) (1b). Each blue line depicts the rate of birth for a particular state of adaptation to the environment, ranging from perfectly adapted *ε*_*μ*_ = 0, as dark blue (top line), to strongly maladapted *ε*_*μ*_ = 2, as light blue (bottom line). The lower panels show the same relationships for cases 2a and 2b. Colored disks show how the 4 scenarios match to figures 3 and 4.

| Environment enters via: | Density Independent |  | Density Dependent |  |
| --- | --- | --- | --- | --- |
| Birth | Case 1a | $b(\gamma_0, N(t))$<br>$d(0, 0)$ | Case 1b | $b(0, \gamma_1 N(t))$<br>$d(0, 0)$ |
| Death | Case 2a | $b(0, 0)$<br>$d(\gamma_0, N(t))$ | Case 2b | $b(0, 0)$<br>$d(0, \gamma_1 N(t))$ |

### Case 2a-2b, dynamic death models

In Case 2a we now incorporate the environmental effect into *d*(*γ*_0_, *γ*_1_) in a density independent way as shown in Figure 1c. In Case 2b, as in Case 2a, the environment alters *d*(*γ*_0_, *γ*_1_), but now alters population response to density as shown in Figure 1d. In both cases holding *b*(*γ*_0_, *γ*_1_) constant.

This yields the four model cases described above and laid out in Table 1.

### Stochastic framework

We used the above ordinary differential equation framework to develop a stochastic simulation algorithm (SSA or birth-death process) using the direct method described by (Gillespie 1977), adapted to allow heritable variation in individual traits. Stochasticity occurs in the model as a result of the random selection of birth and death events (demographic stochasticity), and random mutations during reproduction. This framework is apt for testing our assumptions because true extinctions are possible, and evolution occurs as a result of heritable individual variation that emerges from our assumptions about population demography.

We initialize the model with 35 individuals with traits drawn from a uniform distribution ranging from −1 to 1, the same range as the possible environmental values. Integration of the model starts by first determining the time until the next event, which is randomly sampled from an exponential distribution with mean 1/E, where E is the sum of all possible events (birth or death of each individual):

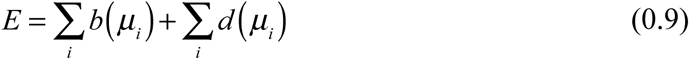

After the current time *t* is updated, the specific event that occurs is determined by randomly choosing among all possible events, weighted according to differences in their rates. For example, the probability that the next event is a death of the ith individual is *d*(*μ*_*i*_) *E*. If an individual dies, it is removed from the population and the entire process is repeated. If an individual reproduces, a random variable on a uniform (0,1) distribution is chosen. If this value is greater than 0.01, the offspring is assigned the parent’s trait value; otherwise, the offspring is given a trait value that is equal to the parent trait value plus a mutation value randomly drawn from a range of −0.3 to 0.3. This sequence of steps mimics mutation-limited evolution in an asexual population. A similar eco-evolutionary framework is described in Delong & Gilbert 2016; however their approach differs slightly from ours because they first aggregate rates of birth and death to the population level, and then randomly assign the individual to experience the event. This results in an underestimate in the response to selection, but leads still to the same equilibrium.

### Simulations

We conducted simulations across a log-linear range of frequencies (*f*) of environmental change. For each frequency of environmental change, we conducted 512 independent replicate simulations. We ran the model for 500 time steps before recording the trait values of each individual, as well as the population size and all simulations continued for another 10000 time steps or until extinction occurred. Trait-environment correlations were computed for the mean phenotype and environment value using Pearson correlation coefficients. To provide a basis of comparison, we also conducted simulations where mutation driven evolution did not occur.

Lastly, we conducted simulations utilizing an environment that changes in a logistic manner

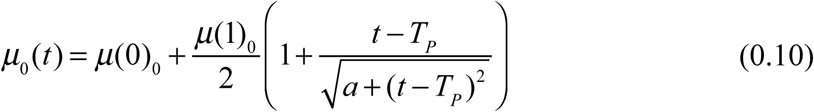

where *μ*_0_ is −1, *μ*_1_ is 1, a is 800 and *T*_*P*_ or the time at which the environment changes is 600. The slope of the environmental change is determined by a, which we chose to be a similar slope of change to our sinusoidal environment *f* = 0.015. We used this additional case to showcase a more traditional type of environmental change to observe evolutionary rescue. Simulations were conducted using Wolfram *Mathematica* v11.0 on a iMac Pro with 18 Xeon W cores.

## Results/Discussion

Our results show that evolutionary rescue is affected when the environment influences different demographic rates and processes. We begin by discussing the resulting extinction dynamics when considering populations that cannot undergo evolution, followed by populations that have the capacity for mutation driven evolution. The four models we consider here are calibrated to produce the same behavior when the environment is held constant; the population will approach an equilibrium density that is determined by the environment, but is consistent across all cases. At equilibrium, however, the turnover rates (approximated by 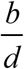) differ among the models in which birth rates vary amongst individuals and those in which death rates vary (see figure 1). Consistent differences also emerge among the models incorporating the density independent and density dependent environmental interaction; particularly at low densities, the effect of trait variation is strongly buffered in the latter cases. These differences give rise to the results depicted in figure 2.

**Figure 2.**
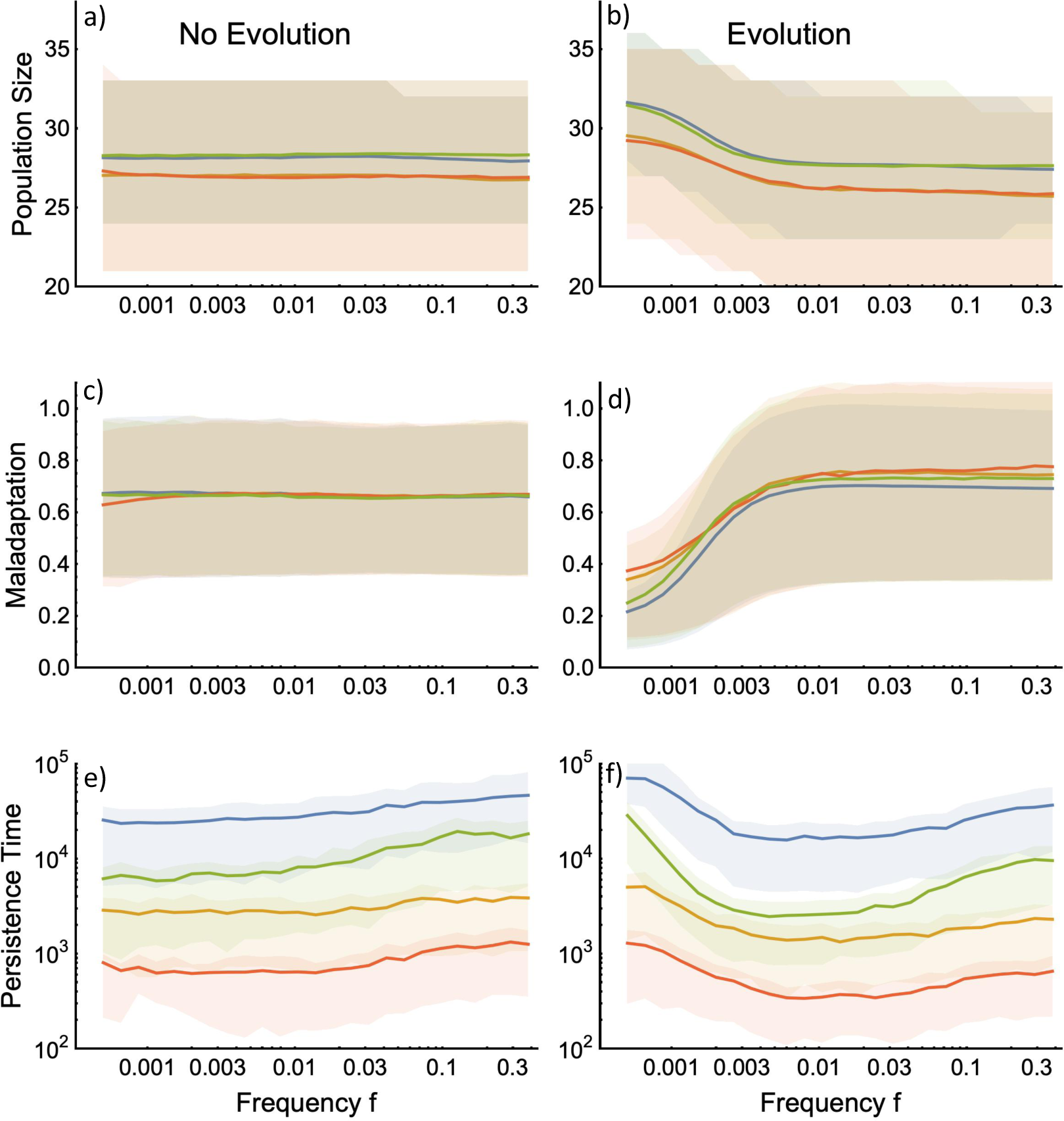
Population dynamics of the four model cases without and with a complete evolutionary dynamic. For population size (a,b) and maladaptation (c,d), the solid lines give the ensemble means of all model replicates and times and the shaded areas show the 25^th^ and 75^th^ percentiles of the distribution. For persistence time (e,f), the solid lines give the means across model replicates and the shaded areas show the 25^th^ and 75^th^ percentiles of the distribution. Maladaptation is measured as the difference between the mean population trait and the environmental value. The blue line represents case 1b, the green line case 1a, the orange line case 2b, and the red line case 2a, as shown in Figure 1.

### Demographic results without evolution

The four models exhibit a consistent ranking of mean persistence time across the entire range of frequencies of environmental change we considered. Mean persistence was greater in populations whose birth rates (rather than death rates) were environmentally influenced, and when the environment affected the strength of density dependence. In the absence of evolution, the most persistent populations were of the form outlined in case 1b, followed by case 1a, where there is a density-environment interaction in the birth rate and where the environment acts on the birth rate independent of density respectively. These were followed by case 2b then 2a the populations where the environment altered the strength of density dependence and acted independent of density on the death rate. This ranking in persistence is easily explained by the ecological differences among the models, considering in particular their behavior when population sizes are small (i.e., as populations are near extinction).

First, populations with birth as the responsive trait persist longer than those with death as the responsive trait due to the greater demographic stochasticity in death models which increases extinction at small pop sizes. The intrinsic growth rate of the population is determined by the difference between the birth and death rate, while demographic stochasticity is determined by the sum of the birth and death rate (Nisbet and Gurney 2003; Palamara et al. 2016). Although our models are parameterized so that they have the same *K*_*A*_ and *K*_*B*_ for when *b* − *d* = 0, the sum of *b* and *d* at these equilibrium points is four times higher in the death models (Case 2a and 2b). Hence the death models have much higher demographic stochasticity than the birth models (Figure 1), and it is clear that demographic stochasticity increases extinction probability at low population sizes (Lande 1993; Melbourne and Hastings 2008). Furthermore, demographic stochasticity increases the variance in population size, as we see in figure 2 (a,b). High fluctuations in vital rates has been shown to decrease population growth due to an increase in variation in the population growth rate (May 1973; Jonsson and Wennergren 2019). Accordingly, various species have been shown to be particularly vulnerable to highly variable adult survival, leading to a higher extinction risk (Lande 1988; Caswell et al. 1999; Jonsson and Ebenman 2001).

Secondly, at low densities, models where the environment interacts with the strength of density dependence maintain higher average (and less variable) population size since maladaptation to the environment has a diminishing impact as population size declines (Fig 1b, 1d). This is reasonable as populations with highly variable growth rates have been shown to be particularly vulnerable to extinction (Leigh 1981; Lande and Orzack 1988). Furthermore it has been shown with a discrete time model that when the environment is a density dependent term it produces a multiplicative effect on population size, and these populations have more strongly bounded populations (Ferguson and Ponciano 2015). As shown in figure 1 (b and d), at low population sizes the density dependent environmental effect has lower variation than the density independent environmental effect, while the opposite is the case at large population sizes. These differences in variation translate into longer persistence times of the models where environmental change alters the affect of density (case 1b, 2b) relative to those where environmental change alters the vital rates independent of density (case 1a, 2a). Although the environmental density effect increases variation at high population sizes, it is favorable when populations are small as they are better able to rebound.

All four scenarios exhibit a rising persistence time as the frequency of environmental variation increases. This is driven by a phenomenon known as “ecological tracking”; when a population ecologically tracks its environment, changes in the environment are re-expressed in the population dynamics as correlated changes in density. Here, where the environment changes sinusoidally, ecological tracking generates population dynamics that exhibit a noisy cycle at the same frequency as the environment (figure 3a,c); however, the tracking response of population diminishes as *f* increases. (May 1976) suggested that the quantity, which represents the system’s dominant eigenvalue, represents a threshold frequency above which tracking does not occur in the Logistic model, but the exact relationship between tracking and the frequency of oscillations is best described as a continuous sigmoid function (Vasseur 2007). The stronger tracking response generated at low frequencies of environmental variation leads to greater variation in population density (both above and below the mean) and thus greater extinction risk. This effect has been shown for a variety of ecological scenarios (Heino et al. 2000; Schwager et al. 2006).

**Figure 3.**
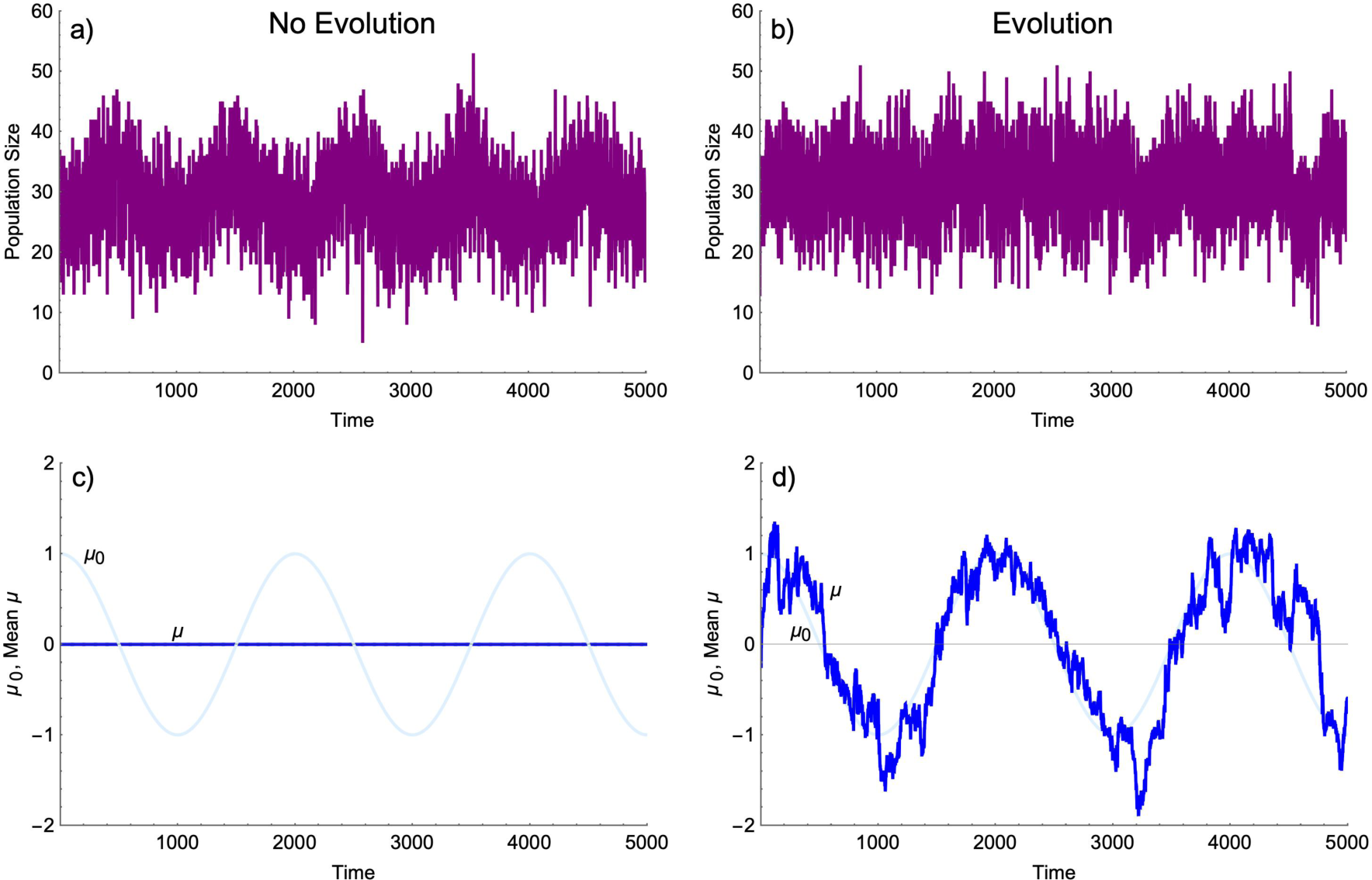
Ecological tracking occurs when the population size (a) exhibits a correlated pattern of variation with the environment (here *μ*_0_) (panels a and c). In this example all individuals have the same trait value, there is no evolution, and *f* = 0.0005. Panels b and d show evolutionary tracking where the mean trait in the population closely follows the environment, thereby dampening the ecological response to the environmental variation.

### Demographic results with evolution

When the full eco-evolutionary dynamics are present in our models, we find that the persistence ranking of models is maintained, however all four models demonstrate a U-shaped (rather than monotonic) relationship between the frequency of environmental change and mean persistence times. This U-shaped relationship arises due to the interplay between ecological and evolutionary tracking of the changing environment. Evolutionary tracking occurs when changes in the environment are slow enough that they can be re-expressed as correlated changes in the mean or modal trait value(s) of the population. Importantly, evolutionary and ecological tracking are interdependent, here forming a link between ecology and evolution. As evolutionary tracking strengthens, ecological tracking is diminished because a population that adapts quickly does not experience the same extent of variation in its vital rates and parameters (here *r* and *K*) (See figure 3b,d). As ecological tracking generally has a negative effect on persistence, evolutionary tracking generates a benefit mitigating the population’s response to ecological tracking. Given the assumptions of our model (mutations per birth, mutation effect size, and population size) evolutionary tracking occurs beginning at approximately *f* = 0.005. Here it can be seen that the deviation between traits and the environmental optimum tends to decline at low frequencies (figure 2d), leading to an increase in the population size and mean persistence times. Together the evolutionary and ecological tracking lead to the U-shaped response to frequency. Variation in population size is not only caused by variation of demographic stochasticity between different vital rates, but also by intraspecific trait variation. Since any individual can give birth in dynamic death models, they have more trait variation in the autocorrelated environments, (low *f*) which increases the effect of maladaptation on their death rate. But as the *f* increases the effect of maladaptation becomes the same across the models.

The eco-evolutionary dynamic that is responsible for an increase in persistence times at low frequencies of environmental fluctuation, also leads to a reduction in persistence time at intermediate and high frequencies (Figure 4). This reduction is due to mutational loading (Higgins and Lynch 2001) which is here exacerbated by the fact that mutations which might be immediately favorable in the population become quickly deleterious as the environment oscillates. This confounding kind of evolution is most likely to occur at intermediate frequencies, where complete evolutionary tracking is unlikely, but random chance allows momentary “misleading” evolutionary changes to occur. We see a slight inflation of the mean and range of maladaptation in our eco-evolutionary models (figure 2c) relative to those without evolution, reinforcing this mechanism. All of our models transition from a detrimental, to a beneficial effect of the eco-evolutionary dynamic near *f* = 0.005. Determining how this threshold relates to the life-history parameters of natural populations will provide important information about the potential for evolution to buffer populations from extinction in oscillating environments. Note that in Figure 2c, the mean line is slightly decreased at low f for the death models. This is due to the higher trait variation exhibited in these models as previously discussed, causing a larger deviation from the optimal trait condition.

**Figure 4.**
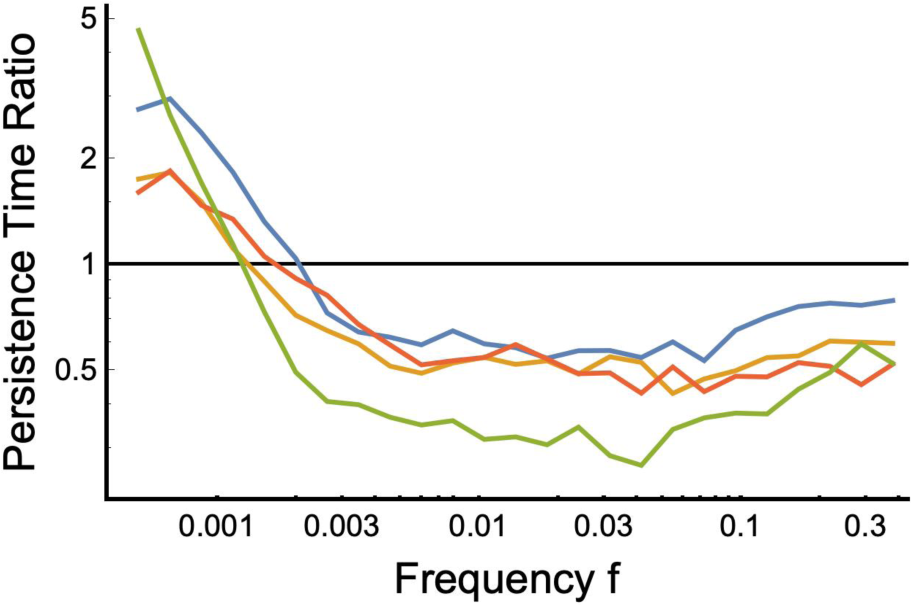
The quotient between the mean persistence time of populations that exhibited evolution and the mean persistence time of populations that did not undergo evolution. For values above one, evolution was beneficial for persistence, for those below one, evolution had a negative impact. Evolutionary tracking increased persistence time for populations when the environmental fluctuation frequency was low.

### Consequences of environmental effects on different demographic rates

In natural populations we see that the demographic rates that are selected upon, and how density dependence responds varies. Some populations may respond to environmental change in a density independent way as in cases 1a, 2a (Dempster 1983; Brewer and Peltzer 2009) while some are likely to show an increase in the intensity of density dependence as in cases 1b, 2b (Aanes et al. 2000; Coulson et al. 2001), with varied key demographic rates, (birth or death). These results emphasize the importance of taking specific demographic parameters into account into our models in the light of evolutionary rescue. Furthermore, these results suggest that environmental change that primarily causes an increase in mortality independent of density will be the most destructive to natural populations (Case 2b). We see dynamics such as this when environmental changes drive populations to physiological limits, natural disasters, severe weather, and pollution. For example, a change in oxygen composition in a marine ecosystem may affect a population regardless of density (Brewer and Peltzer 2009), or an increase in heavy metal contamination may similarly increase mortality regardless of population size (Santala and Ryser 2009).

According to our results the populations that will benefit the most from evolutionary rescue will be those whose fecundity responds to an environmental change in a density dependent way. This may be exemplified in cases where the availability of, or access to resources diminishes or changes with environmental change. This leads to the malnutrition and lower fecundity of some individuals (Jaumann and Snell-Rood 2019) but importantly in this case, as the population size declines the effect of the environmental stress weakens. Note that density dependence can also decrease due to environmental change in areas where the change is favorable (take the case of invasive species and pests), further increasing persistence potential (Ouyang et al. 2014). From these results we recommend that long-term studies incorporate fine demographic data when feasible. Further analysis should be done to fine tune the relevant parameters that play a role in evolutionary rescue, so that we may one day be able to predict and promote evolutionary rescue in the wild.

### Consequences of our model assumptions

Our modeling framework assumes asexual reproduction and a link between the environment and demographic parameter that is controlled by a single trait. Most empirical and theoretical work suggests that sexual recombination leads to an increased rate of evolution, as it is beneficial when mutations are common and have a small effect size (Crow and Kimura 1965). Recombination can also pose the opposite effect by allowing maladaptive traits to persist longer in the population, leading to a greater genetic load on population fitness. Incorporating recombination to assess any differences in outcome will surely be relevant given the diversity of mating systems in nature. Furthermore, singular step mutations are what allow the population as a whole to track the changing environment, as opposed to a genotype phenotype mapping that is not one to one. This may be representative of populations with a narrow genetic basis for which adaptation to the environment can occur, such as what has commonly been seen in drug resistance (MacLean et al. 2010). That being said, in nature some cases of environmental change will surely require multiple traits to evolve for the population to persist. The utility of this model though is that it is comparative, it is likely we will see the same trends in a multi-trait model but this will surely be fruitful to investigate as we bring our models towards realism. This will become even more relevant with the incorporation of species interactions. Competition can both inhibit and promote evolutionary rescue in different cases (Osmond and de Mazancourt 2013) and has shown to be a relevant component in the study of population persistence.

Lastly, the environment in this model lacks environmental stochasticity, which has been shown to play a role in the potential for populations to evolve to track the changing environment (Ovaskainen and Meerson 2010; Fey and Wieczynski 2017). But, because we utilize a fluctuating environment instead of the single step change commonly utilized in evolutionary rescue studies, we are able to characterize the ability for a population to continuously adapt to a changing environment. In this way we are able to see populations undergoing evolutionary rescue again and again, in order to better understand the mechanisms underlying this dynamic. In environments undergoing non-cyclic changes, the rate and extent of environmental change together form a critical axis on which the success of evolutionary rescue (or more appropriately eco-evolutionary rescue) can be measured. Generally, the potential for eco-evolutionary rescue is assessed using a singular environmental change, e.g. from low to high concentrations of salt, or cold to warm temperatures, (Doebeli and Dieckmann 2003; Crump et al. 2004; McCain and Grytnes 2010) and the typical pattern of population and trait dynamics are easily explained using the concepts of ecological and evolutionary tracking applied above; when traits are able to track the environmental change quickly enough, ecological changes are dampened enough to prevent extinction. Thus, our model, which incorporates a cyclic environmental change, is a useful predictor of how different assumptions about life history will alter the propensity of eco-evolutionary rescue. We confirm that our results are not an outcome of this cyclic environment, as the same persistence ranking results from a sinusoidal shift in the environment (Figure 5).

**Figure 5.**
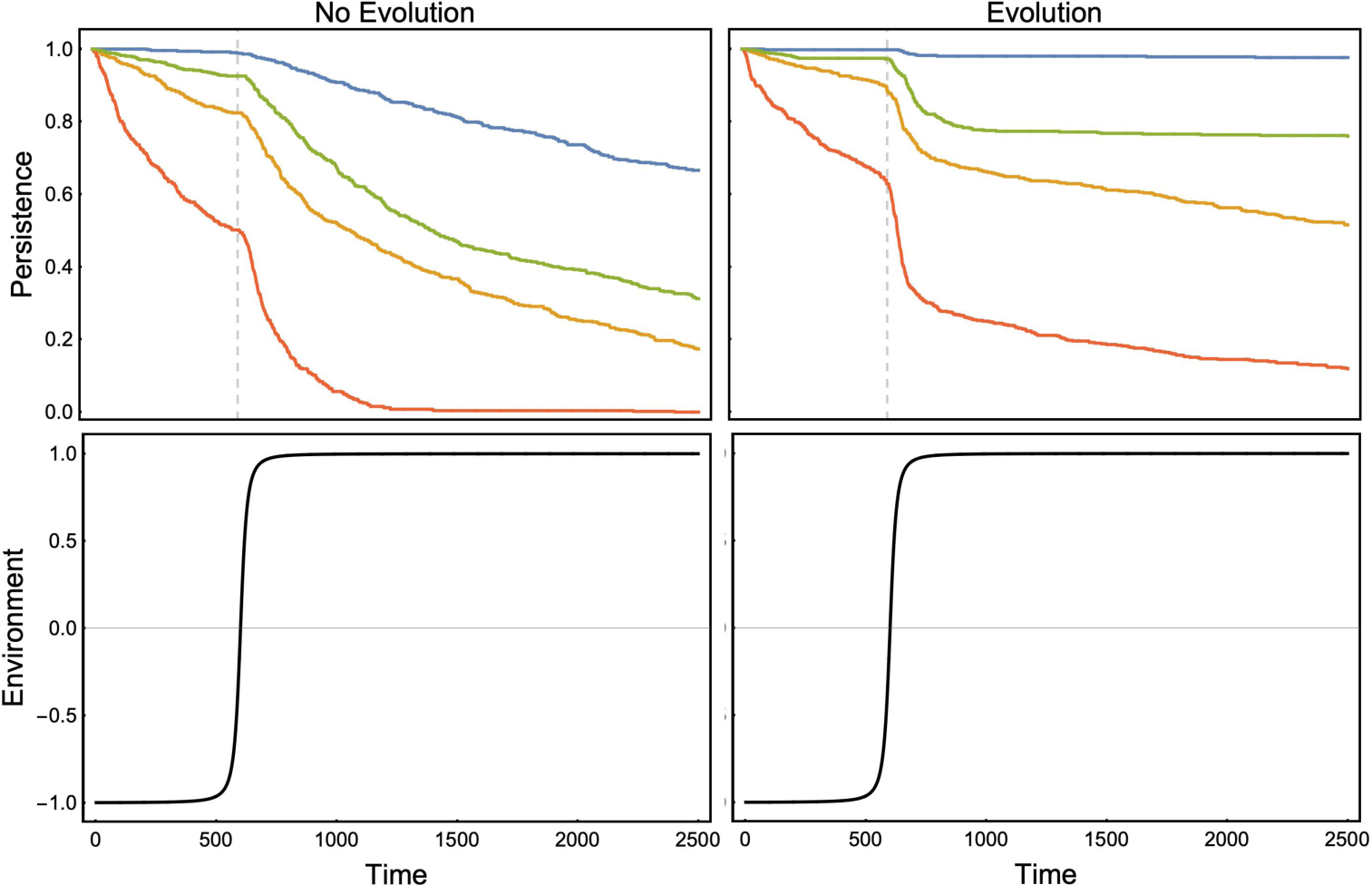
The proportion of persisting populations over time. These plots portray a typical evolutionary rescue scenario with a sigmoidal environment. The top panels depict the proportion of surviving populations over time out of 512 replicates for case 1a (green), 1b (blue), 2a (red), and 2b (yellow).

The study of evolutionary rescue has increased notably in the past decade, and although we have elucidated a reduced set of relevant factors, the interplay between demography and evolutionary rescue is still largely unknown. We show that models with varied dynamic demographic parameters with the same carrying capacities and initial conditions have different probabilities of undergoing evolutionary rescue following environmental change. Therefore, comparative evolutionary demography provides a lens with which we can understand how different populations may be more or less likely to persist alongside environmental change. As emphasized in previous studies, evolutionary rescue in these models occurs when the rate of environmental change, or the fluctuation frequency is slow enough for the population to evolutionarily track the changing trait optimum as shown in figure 3b,d (Perron et al. 2008; Lindsey et al. 2013). Although the current model does not take into account spatially heterogeneous environments or interspecific competition, it provides a starting point to better understand the interplay between evolutionary demography and evolution to a changing environment. We find that changing the demographic parameter that selection acts on, as well as the way in which selection alters density dependence, changes a populations propensity to avoid extinction via evolutionary rescue.

## Conclusion

We show that when evolution is occurring in a system, the extinction probabilities vary given different dynamic demographic parameters. This work is the first to show that populations whose abundance is determined by changes in different key demographic rates have different probabilities to avoid climate-induced extinction via evolution. This comes into play in how well a population can evolve to have high fitness in a changing environment, and the ability of a population to rebound from small population sizes. Our findings show the importance of explicitly incorporating environmental change and density dependence into equations describing population demographic rates. In our study the environment provides the selective pressure on individuals, and unlike in previous work the shape of this selective pressure is shown to differ between commonly used models. This result would not have been shown had we focused on a purely ecological or evolutionary model, this interplay is what allows us to make novel insights into if and how population persistence will be altered by climate change. Furthermore, incorporating selection and trait evolution into models on ecological time scales is an important research priority. This work shows that natural populations that have different key demographic rates will likely respond differently to climate change, and this information should be explicitly incorporated into models that predict extinction due to climate change.

In order to minimize extinction of natural populations alongside changing environmental conditions such as climate change, we must be able to make decisions without complete data describing future phenomena. It is therefore vital to create theory that can aid scientists and wildlife managers alike in understanding how natural populations respond to escalating rates of environmental challenge. This includes techniques utilizing the population data we already have, to use the past as a proxy for the future, as well as techniques utilizing our understanding of evolution to form ideas of how populations can adapt and how we can help them to adapt to persist into the future. Our current lack of understanding of the combined effect of ecological and evolutionary dynamics on the outcome of climate change, poses a challenge to produce theoretical and experimental work investigating these mechanisms. Already scientists are corroborating theoretical hypotheses with experimental results for concepts such as rate of environmental change, initial population size, and genetic variability (Bell and Gonzalez 2009, 2011; Martin et al. 2013). The results provided in this study provide us with new testable hypotheses that we can test utilizing experimental evolution. The comparative framework we’ve established allows us to test the probability of population rebound post decline due to environmental change between populations whose demography responds differently.

It is clear that in order to asses a populations propensity for evolutionary rescue, we must pay attention to the specific life history parameters that determine population size both with and without environmental change. That is, what is the key factor that determines population size, what role density dependence plays and how environmental change alters the vital rates and their response to density (Coulson et al. 2008). The way that the environment alters population vital rates and response to density in predictive models is often simplified in the literature when using data driven frameworks that predict population size based on current habitat (Guisan and Zimmermann 2000; VanDerWal et al. 2009; Thuiller et al. 2014), and theoretical frameworks where the environmental change acts on r directly (Vasseur et al. 2011; Cropp and Norbury 2019). This can conceal population response to environmental change as r is an aggregate of many processes (fecundity, mortality, dispersal etc) that may respond differently to environmental change.

